# Template-based prediction of vigilance fluctuations in resting-state fMRI

**DOI:** 10.1101/234179

**Authors:** Maryam Falahpour, Catie Chang, Chi Wah Wong, Thomas T. Liu

## Abstract

Changes in vigilance or alertness during a typical resting state fMRI scan are inevitable and have been found to affect measures of functional brain connectivity. Since it is not often feasible to monitor vigilance with EEG during fMRI scans, it would be of great value to have methods for estimating vigilance levels from fMRI data alone. A recent study, conducted in macaque monkeys, proposed a template-based approach for fMRI-based estimation of vigilance fluctuations. Here, we use simultaneously acquired EEG/fMRI data to investigate whether the same template-based approach can be employed to estimate vigilance fluctuations of awake humans across different resting-state conditions. We first demonstrate that the spatial pattern of correlations between EEG-defined vigilance and fMRI in our data is consistent with the previous literature. Notably, however, we observed a significant difference between the eyes-closed (EC) and eyes-open (EO) conditions finding stronger negative correlations with vigilance in regions forming the default mode network and higher positive correlations in thalamus and insula in the EC condition when compared to the EO condition. Taking these correlation maps as “templates” for vigilance estimation, we found that the template-based approach produced fMRI-based vigilance estimates that were significantly correlated with EEG-based vigilance measures, indicating its generalizability from macaques to humans. We also demonstrate that the performance of this method was related to the overall amount of variability in a subject’s vigilance state, and that the template-based approach outperformed the use of the global signal as a vigilance estimator. In addition, we show that the template-based approach can be used to estimate the variability across scans in the amplitude of the vigilance fluctuations. We discuss the benefits and tradeoffs of using the template-based approach in future fMRI studies.

## 1. Introduction

In recent years, there has been a growing interest in using resting state functional magnetic resonance imaging (fMRI) as a tool to study the functional organization of the brain. Since resting state fMRI signals contain a mixture of neural and non-neural components, the ability to gain insight into human brain function from resting state data critically relies on the ability to understand and account for major sources of the variability in the data. Retaining a constant level of vigilance or wakefulness is difficult or perhaps impossible, especially during resting state scans, due to the absence of an engaging task. In fact, a recent study revealed that about a third of participants lost wakefulness within the first 3 minutes of a resting-state scan and that half of the participants lost wakefulness after 10 minutes (Tagliazucchi and Laufs, 2014). Given that there are many neural and neurotransmitter systems involved in vigilance state changes (Oken et al., 2006), fMRI signal changes relating to vigilance modulation can potentially mask the ability to detect other neural signals of interest during an fMRI experiment. For example, prior work has shown that functional connectivity is altered during the loss of wakefulness (Haimovici et al., 2017; Laumann et al., 2017), after taking caffeine (Wong et al., 2013), and by changes in state between the eyes open and closed conditions (Yan et al., 2009). Yet, a broad range of fMRI resting state studies typically make the implicit assumption that all participants are in similar states of wakefulness or vigilance.

While EEG can be used to monitor alertness and vigilance in real-time (Jung et al., 1997), collecting EEG data during fMRI experiments is usually not practical. Therefore, it becomes important to have methods for estimating vigilance fluctuations using the fMRI data alone. Toward this goal, (Tagliazucchi et al., 2012; Altmann et al., 2016) demonstrated approaches for inferring discrete sleep stages from windowed fMRI connectivity matrices, and recently, (Chang et al., 2016) showed that continuous variations in alertness between wakefulness and the transition to light sleep may be tracked using an fMRI template-based approach in awake monkeys sitting in complete darkness.

The primary aim of the present study is to determine whether this template-based approach can successfully predict vigilance fluctuations in typical resting state scans conducted in humans. We used simultaneously acquired EEG-fMRI data to derive the vigilance template and evaluate the approach in healthy volunteers in both the eyes-closed (EC) and eyes-open (EO) conditions. For both conditions, data were also acquired before and after the administration of a caffeine dose. We examined the differences between the templates (i.e., EEG-fMRI correlations) derived from different conditions and evaluated how well these templates can predict both the time course of vigilance fluctuations and the overall amount of temporal variations in the vigilance state (i.e. standard deviation of vigilance fluctuations). Finally we compared the template-based approach to the use of the global signal as a vigilance predictor, motivated by prior work (Falahpour et al., 2016) showing significant correlations between the global signal and vigilance fluctuations.

## 2. Methods

### 2.1. Experimental protocol

In this work we used data from the protocol described in our prior work (Wong et al., 2013, 2015). Details of the experimental protocol, acquisition and fMRI/EEG data pre-processing were described in those articles and are repeated here for the reader’s convenience. Twelve healthy volunteers were initially enrolled in this study after providing informed consent. Two subjects were not able to complete the entire study, resulting in a final sample size of 10 subjects (4 males and 6 females, aged 24-33 years, with an average age of 25.6 years). As prior work has shown that differences in dietary caffeine consumption may cause variability in the BOLD response (Laurienti et al., 2002), we recruited caffeine-naive subjects who consumed less than 50 mg caffeine daily (as assessed over a two month period prior to the study).

A repeated measures design was used, with each subject participating in two imaging sessions: a caffeine session and a control session. The order of the two sessions was randomized in a double-blinded manner. For each session, the operator obtained a capsule that contained 200 mg of either caffeine or cornstarch. The two imaging sessions were separated by at least two weeks. Each session consisted of a pre-dose and a post-dose imaging section, with each section lasting for about one hour. Upon completion of the pre-dose section, participants were taken out of the magnet and given the capsule. The subject was then placed back in the scanner, with the first functional scan of the post-dose section obtained approximately 40 min after capsule ingestion. This interval was chosen based on studies showing that the absorption of caffeine from the gastrointestinal tract reaches 99% about 45 min after ingestion with a half-life ranging from 2.5 to 4.5h (Fredholm et al., 1999).

Each scan section consisted of (1) a high-resolution anatomical scan, (2) arterial spin labeling scans (these results are reported in our prior study (Wong et al., 2012) but not considered here), and (3) two 5 minute resting-state scans with simultaneous EEG recording (one eyes-closed and one eyes-open). Subjects were instructed to lie still in the scanner and not fall asleep during resting-state scans. The order of the eyes-open and eyes-closed scans was randomized. During the eyes-open resting-state scans, subjects were asked to maintain attention on a black square located at the center of a gray background. During the eyes-closed resting-state scans, subjects were asked to imagine the black square.

An impedance check was performed prior to each EEG data acquisition while the subject was in the desired state (EC or EO) for at least 1.5 min. EEG data acquisition began 30 seconds before each fMRI scan and ended 30 seconds after. Thus, the subject was in the desired state (EC or EO) for at least 2 minutes prior to the acquisition of the combined EEG and fMRI data. Field maps were acquired to correct for magnetic field inhomogeneities.

### 2.2. MR data acquisition

Imaging data were acquired on a 3 Tesla GE Discovery MR750 whole body system using an eight-channel receive coil. High resolution anatomical data were collected using a magnetization prepared 3D fast spoiled gradient (FSPGR) sequence (TI = 600 ms, TE = 3.1 ms, flip angle = 8°, slice thickness = 1 mm, FOV = 25.6 cm, matrix size =256 × 256 × 176). Whole brain BOLD resting-state data were acquired over 30 axial slices using an echo planar imaging (EPI) sequence (flip angle = 70°, slice thickness = 4 mm, slice gap = 1 mm, FOV = 24 cm, TE = 30 ms, TR = 1.8 s, matrix size= 64 × 64 × 30). Field maps were acquired using a gradient recalled acquisition in steady state (GRASS) sequence (TE1 = 6.5 ms, TE2 = 8.5 ms), with the same in-plane parameters and slice coverage as the BOLD resting state scans. The phase difference between the two echoes was then used for magnetic field inhomogeneity correction of the BOLD data. Cardiac pulse and respiratory effect data were monitored using a pulse oximeter (InVivo) and a respiratory effort transducer (BIOPAC), respectively. The pulse oximeter was placed on each subject’s right index finger while the respiratory effort belt was placed around each subject’s abdomen. Physiological data were sampled at 40 Hz using a multi-channel data acquisition board (National Instruments).

### 2.3. EEG data acquisition

EEG data were recorded using a 64 channel MR-compatible EEG system (Brain Products, Munich, Germany). The system consisted of two 32 channel Brain Amp MR Plus amplifiers powered by a rechargeable battery unit. The system was placed behind the scanner bore, which was connected using a 125 cm long data cable to a Brain Cap MR with 64 recording electrodes (Brain Products, Munich, Germany). All electrodes in the cap had sintered Ag/AgCl sensors incorporating 5 kΩ safety resistors. The separate ECG electrode had a built-in 15 kΩ resistor. The arrangement of the electrodes in the cap conformed to the international 10/20 standard. FCz and AFz were the reference and ground electrodes, respectively. The EEG data were recorded at a 5 kHz sampling rate with a passband of 0.1-250 Hz. A phase locking device (Syncbox, Brain Products, Munich, Germany) was used to synchronize the clock of the EEG system with the master clock of the MRI system. Before each scan section, the electrode impedances were set below 20 kΩ, while the impedances of the reference and ground electrodes were set below 10 kΩ. Prior to recording EEG data in each resting-state scan, a snapshot of the electrode impedance values was taken from the computer screen. One EEG dataset was created for each 5-min resting-state scan.

### 2.4. MR data processing

AFNI and FSL were used for MRI data pre-processing (Cox, 1996; Smith et al., 2004; Woolrich et al., 2009). The high resolution anatomical data were skull stripped and segmentation was applied to estimate white matter (WM), gray matter (GM) and cerebral spinal fluid (CSF) partial volume fractions. In each scan section, the anatomical volume was aligned to the middle functional volume of the first resting state run using AFNI. The anatomical volume in the post-dose scan section was then registered to the pre-dose anatomical volume, and the rotation and shift parameters obtained from this registration were applied to the post-dose functional images. The first 6 TRs (10.8 s) of the BOLD data were discarded to allow magnetization to reach a steady state. A binary brain mask was created using the skull-stripped anatomical data. For each slice, the mask was eroded by two voxels along the border to eliminate voxels at the edge of the brain (Rack-Gomer and Liu, 2012). For each run, nuisance terms were removed from the resting-state BOLD time series through multiple linear regression. These nuisance regressors included: i) linear and quadratic trends, ii) six motion parameters estimated during image co-registration and their first derivatives, iii) RETROICOR (2nd order Fourier series) (Glover et al., 2000) and RVHRCOR (Chang et al., 2009; Chang and Glover, 2009) physiological noise terms calculated from the cardiac and respiratory signals, and iv) the mean BOLD signals calculated from WM and CSF regions and their respective first derivatives, where these regions were defined using partial volume thresholds of 0.99 for each tissue type and morphological erosion of two voxels in each direction to minimize the inclusion of gray matter.

### 2.5. EEG data processing

#### 2.5.1. Removing the artifacts

Vision Analyzer 2.0.1 software (Brain Products, Munich, Germany) was used for MR gradient removal using an average pulse artifact template procedure (Allen et al., 2000). An average template (over all volumes) was created using the volume-start markers from the MRI system and then subtracted from the individual artifacts. After gradient artifact removal, a low pass filter with a cutoff frequency of 30 Hz was applied to all channels and the processed signals were down-sampled to 250 Hz. Heart beat event markers were created within the Analyzer software.

The corrected data and event markers were exported to Matlab 7 (Mathworks, Inc.), and EEGLAB (version 9) was used for further pre-processing (Delorme and Makeig, 2004). For each EEG dataset, the ballistocardiogram (BCG) and residual artifacts were removed using a combined optimal basis set and independent component analysis approach (OBS-ICA) (Debener et al., 2007; Niazy et al., 2005). To remove the BCG artifact, the continuous EEG data were divided into epochs based on the heart beat event markers. The epochs were stacked into a matrix configuration and a BCG template was created using the first three principal components calculated from the matrix (Debener et al., 2007). The BCG template was then fitted in a least-squares manner and subtracted from each epoch of the EEG data.

After BCG artifact removal, channels that exhibited high impedance values (>20 kΩ) or were contaminated with high levels of residual gradient artifact were identified and discarded from further processing. The impedance values were identified using the snapshot of channel impedances acquired before the beginning of each scan. To identify channels contaminated with residual gradient artifact that were not adequately removed using the Analyzer software, the continuous EEG data in each channel were bandpass filtered from 15.5 to 17.5 Hz. This frequency band contained the first harmonic of the gradient artifact centered at 16.7 Hz (slice markers were separated by 60 ms). The root mean square (rms) of the filtered time course was calculated for each channel. A channel was identified as a contaminated channel if the rms value was larger than the median plus 6 times the inter-quartile range (Devore and Farnum, 2005), calculated across all channels except the ECG. On average, each channel was included in 95% of the runs in both the eyes-closed and eyes-open conditions (median = 98%, s.d. = 6%). For further analysis, only the included channels for each run were used.

Extended infomax ICA was then performed. Independent components (ICs) corresponding to residual BCG or eye blinking artifacts were identified by correlating all IC topographies with artifact template topographies and extracting the ICs with spatial correlation values of 0.8 or more (Debener et al., 2007; Viola et al., 2009). The corrected data were then created by projecting out the artifactual ICs (Delorme and Makeig, 2004).

As bulk head motion creates high amplitude distortion in the EEG data acquired in the MRI environment, it is desirable to discard the distorted time segments (Jansen et al., 2012; Laufs et al., 2008). To identify the contaminated time segments, the EEG data were bandpass filtered from 1 to 15 Hz (to avoid the first harmonic of the residual gradient artifact centered at 16.7 Hz). A mean amplitude time series was calculated by taking the rms across channels. Outlier detection was performed on the mean time series and the outlier threshold was calculated as the sum of the median value and 6 times the inter-quartile range across time (Devore and Farnum, 2005). Data points with values larger than this threshold were regarded as segments contaminated by motion. Contaminated segments less than 5 seconds apart were merged to form larger segments that were then excluded. A binary time series was created by assigning a 1 to the time points within the bad time segments and a 0 to the remaining good time points. In a recent study, (Jansen et al., 2012) found that predictors derived from an EEG motion artifact were strongly correlated with the BOLD signal when the EEG predictors were convolved with a hemodynamic response function. To indicate the time points reflecting the BOLD response to the motion artifact, we convolved the binary time series with a hemodynamic response function (Buxton et al., 2004) and binarized the resulting output with a threshold of zero to form a second binary time series. Since both EEG and fMRI measures were considered in our study, a final binary time series was created by combining the two binary time series using an OR operation. The binary time series were down-sampled to match the sampling frequency of the BOLD time courses. In the eyes-closed condition, an average of 81% of the time points of the binary time series were indicated as good (i.e. with a value of zero; median = 85%, s.d. = 13%). In the eyes-open condition, the corresponding average was 86% (median = 88%, s.d. = 9%).

#### 2.5.2. Forming the spectrogram

For each EEG channel, a spectrogram was created using a short-time Fourier transform with a 4-term Blackman-Harris window (1311 points, 5.24 s temporal width, and 65.7% overlap). The output temporal resolution of the spectrogram was 1.8 s (i.e. equivalent to the TR of the BOLD resting-state scans). The time points in the EEG spectrogram and BOLD time series that were indicated as potentially contaminated by motion (i.e. values of 1 in the binary time series) were discarded from further analysis.

#### 2.5.3. Sleep staging

Manual sleep staging was performed by a registered polysomnographic technologist using AASM guidelines (AASM, 2009). Specifically, the corrected data (using channels F3, F4, C3, C4, O1 and O2) were segmented into 30 second epochs (with 10 epochs per run) and a sleep stage was scored for each epoch. Based on the sleep staging, it was determined that one subject was not awake during the majority of their scans, and this subject was excluded from further analysis. All of the remaining nine subjects were awake during all 20 epochs of the eyes-open runs. For the eyes-closed runs, five subjects were awake during all 20 epochs. The remaining 4 subjects were awake during the majority of the epochs, but had some epochs with stage N1, with two subjects also exhibiting one or two epochs of stage N2. (See (Wong et al., 2015) for more details.)

## 3. fMRI and EEG metrics

### 3.1. EEG vigilance time series

As described above, a spectrogram was calculated for each EEG channel with the same temporal resolution as the BOLD time series. A global spectrogram was created by computing the rms value (across all channels) at each frequency. Then at each time point, a relative amplitude spectrum was computed by normalizing the spectrum by its overall rms amplitude (square root of sum of squares across all frequency bins). Relative EEG amplitude time series were then computed as the rms amplitude in the following frequency bands (delta and theta: 1–7 Hz, alpha: 7–13 Hz). The same range of frequencies were used for both EC and EO scans (Barry et al., 2007; Wong et al., 2015). Vigilance was defined as the ratio of power in alpha band over the power in delta and theta band (Horovitz et al., 2008; Olbrich et al., 2009; Wong et al., 2013), which is equivalent to the alpha slow-wave index (ASI) (Jobert et al., 1994; Larson-Prior et al., 2009; Müller et al., 2006). Prior studies have shown that decreases in vigilance or wakefulness are characterized by increases in delta and theta band power and decreases in alpha band power (Klimesch, 1999; Strijkstra et al., 2003; Müller et al., 2006), and that the ratio of powers in the high and low frequency bands is correlated with various aspects of BOLD fMRI activity (Horovitz et al., 2008; Olbrich et al., 2009; Wong et al., 2013). In order to form the vigilance time series, rms amplitude in the alpha band was divided by the rms amplitudes in the delta and theta bands at each time point. Vigilance time series were then convolved with the hemodynamic response function to account for the hemodynamic delay. Finally, vigilance amplitude (*aVig*) was computed for each run as the standard deviation of the vigilance time series after removing the bad time points.

### 3.2. fMRI global signal (GS)

For each voxel, a normalized BOLD time series was obtained by subtracting the mean value and then dividing the resulting difference by the mean value. The global signal was formed by averaging the normalized time series across all brain voxels. Correlation between the global signal and vigilance time series was then computed after excluding the bad time points.

## 4. Correlation between vigilance time series and BOLD fMRI data

For each run, the temporal correlation between the EEG vigilance time series and each voxel’s BOLD time series was calculated after excluding the bad time points. Following the approach of Chang et al. (2016), the individual correlation maps will be referred to as individual templates (*Temp_i_*) throughout this work and were derived for all conditions. The definition of the conditions used throughout this work are as follows: ECnonCaf and EOnonCaf refer to all non-caffeine runs (from both the control and caffeine imaging sessions) with eyes-closed and eyes-open, respectively; ECpreCaf and EOpreCaf refer to the runs before caffeine intake that were acquired during the caffeine-related imaging sessions (note these are a subset of the non-caffeine scans); and ECpostCaf and EOpostCaf refer to the runs after caffeine consumption. In order to simplify the presentation, when the meaning is clear from the context, we will also use the terms EC or EO alone to indicate the state of the eyes within a condition. For each non-caffeine condition (ECnonCaf or EOnonCaf) the average of the correlation maps from all runs within that condition excluding one run was used as the Leave-One-Out template (*Temp_LOO_*). Then at each time point, the spatial correlation between the *Temp_LOO_* and the fMRI data of the excluded scan was computed generating a new time series which will be considered as the vigilance estimate (*Vig_est_*). The temporal correlation between the estimate (*Vig_est_*) and the EEG-based vigilance time course (Vig) of the excluded scan was calculated and used as a measure of “predictivity”. The same procedure was repeated for all sessions and runs to obtain the average performance across scans. Additional details regarding the method are presented in (Chang et al., 2016).

### 4.1. Difference in the correlation maps derived from different conditions

Fisher-z transformation was used to normalize the individual correlation maps. Two-sample paired t-tests were then used to assess the group differences across conditions, i.e ECnonCaf versus EOnonCaf, ECpreCaf versus ECpostCaf, and EOpreCaf versus EOpostCaf. We used the modified 3dttest++ (with Clustsim option) from AFNI which uses t-test randomization to minimize the FPR.

### 4.2. Predictivity across different conditions

Predictivity across conditions was also examined by deriving an average template (*Temp_ave_*) from one condition (e.g. ECnonCaf) and then using it to estimate the vigilance fluctuations from another condition (e.g. EOnonCaf).

## 5. Results

For each run, an estimated vigilance time series was found by calculating the spatial correlation between the *Temp_LOO_* and the fMRI data at each time point. The correlations between the estimated vigilance (*Vig_est_*) and the EEG-based vigilance time courses for the non-caffeine sessions (ECnonCaf and EOnonCaf) are shown in Figure 1. Correlation values significantly different from zero (*p* < 0.05) are marked with black asterisks. We used a one-sample t-test on the Fisher z-transformed correlation coefficients to confirm the overall positive predictivity values and Cohen’s d to assess the effect size in both EC (*t*(26) = 7.8, *p* < 10^−6^, *d* = 1.5) and EO conditions (*t*(26) = 12.1, *p* < 10^−6^, *d* = 2.3). No significant difference was found between the EC and EO values (EO-EC: *t*(26) = 1.2, *p* = 0.2, *d* = 0.2).

**Figure 1:**
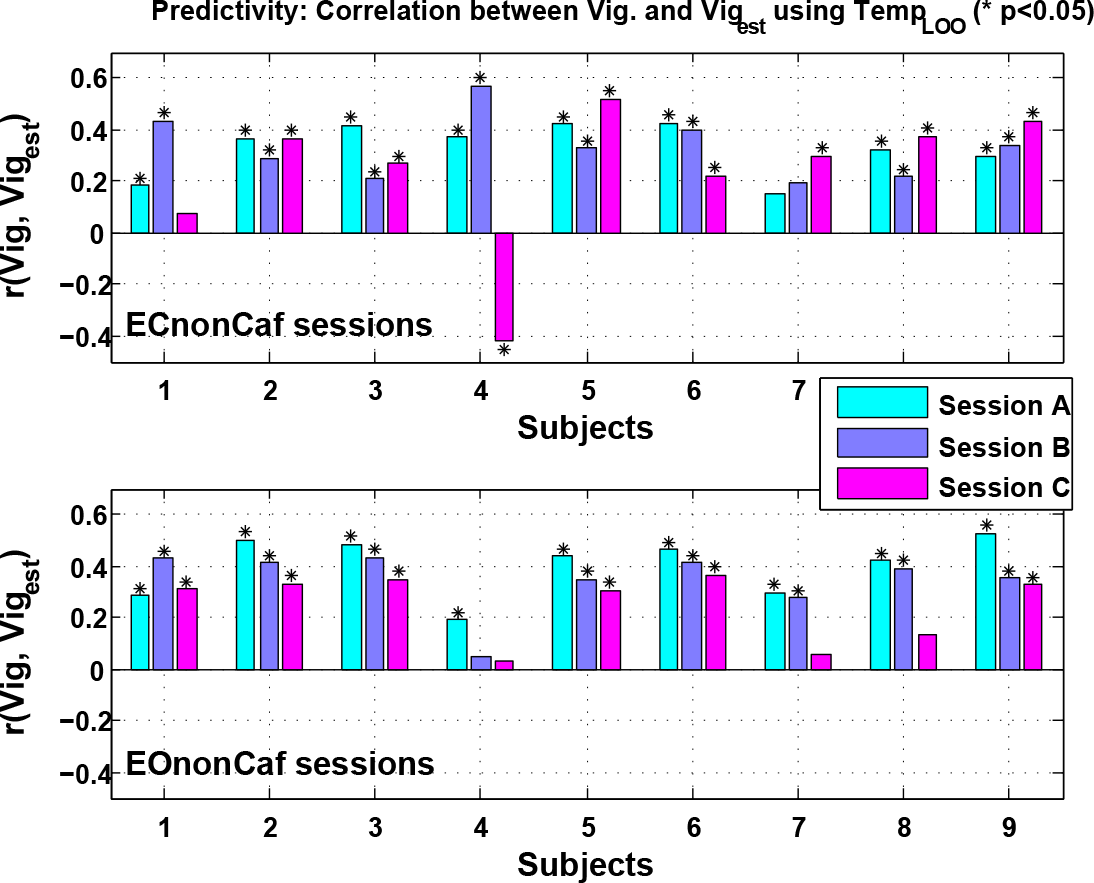
Correlation between the EEG-based vigilance time series and the estimated vigilance using *Temp LOO* across subjects and runs for the ECnonCaf scans on the top row and the EOnonCaf scans in the bottom row. (* p < 0.05)

The average correlation maps, i.e. *Temp_ave_*, derived from the ECnonCaf, EOnonCaf, ECpostCaf, and EOpostCaf conditions and the spatial correlations between them are illustrated in Figure 2. We found that the non-caffeine EC and EO templates were highly correlated (*r* = 0.75, *p* < 10^−6^), and the ECpostCaf template had the lowest correlation with the other templates (*r*(*ECpostCaf, ECnonCaf*) = 0.38, *p* < 10^−6^, *r*(*ECpostCaf, EOnonCaf*) = 0.39, *p* < 10^−6^, *r*(*ECpostCaf, EOpostCaf*) = 0.25, *p* < 10^−6^).

**Figure 2:**
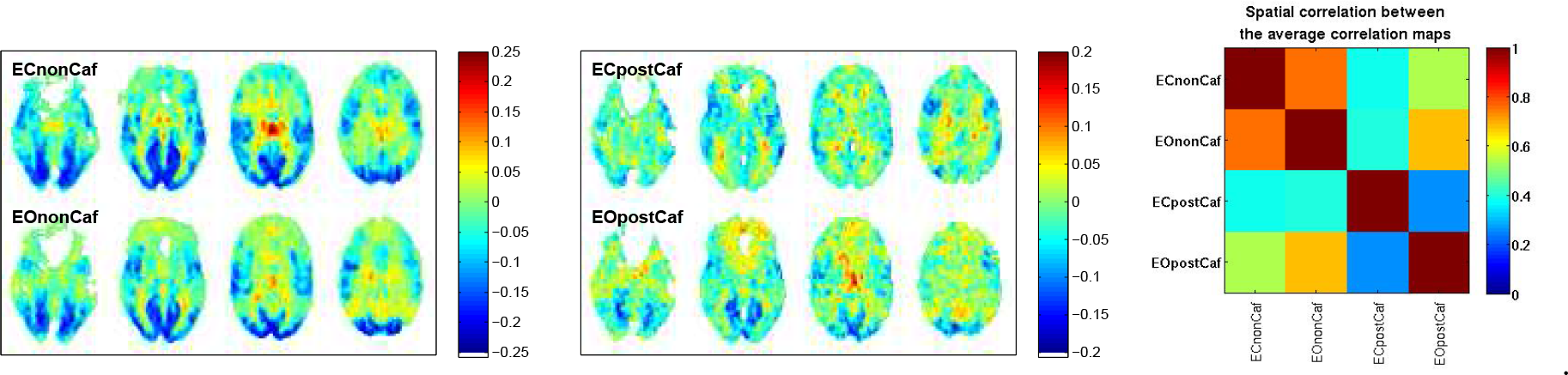
Left: Average of the correlation maps derived from all non caffeine scans, with EC on top and EO in bottom row; Middle: Average of the correlation maps derived from post caffeine session, with EC on top and EO in bottom row; Right: Spatial correlation between the average correlation maps derived from different conditions.

### 5.1. Differences in the correlation maps

Group differences in the vigilance correlation maps derived from the ECnonCaf and EOnoncaf conditions (EC minus EO) are shown in the top row of Figure 3. Maps were first thresholded at *p* < 0.005 voxel-wise, and then thresholded at a cluster size of 69 voxels to correct for multiple comparisons. We found that regions forming the well-known default mode network (posterior cingulate cortex, temporal gyrus, and medial frontal gyrus) showed significant (*p* < 0.05, corrected) reductions in the correlation between the BOLD time series and vigilance in the EC condition as compared to EO, whereas thalamus, cingulate gyrus and insula showed significantly higher correlations in the EC condition. We also examined the effect of caffeine on the correlation maps in both eyes-closed and eyes-open conditions, and the areas showing significant differences (*p* < 0.05, uncorrected) are displayed in the 2nd and the 3rd rows of Figure 3, respectively. None of the significant areas survived multiple comparisons correction but the maps are presented for qualitative comparison between the conditions. We found that caffeine has a stronger effect in the EC condition with significant voxel-wise differences in the posterior cingulate, cuneus, thalamus and right insula. In the EO condition, only a few small clusters in cuneus, preceuneus and left superior pariatal lobule showed significant voxel-wise changes after caffeine.

**Figure 3:**
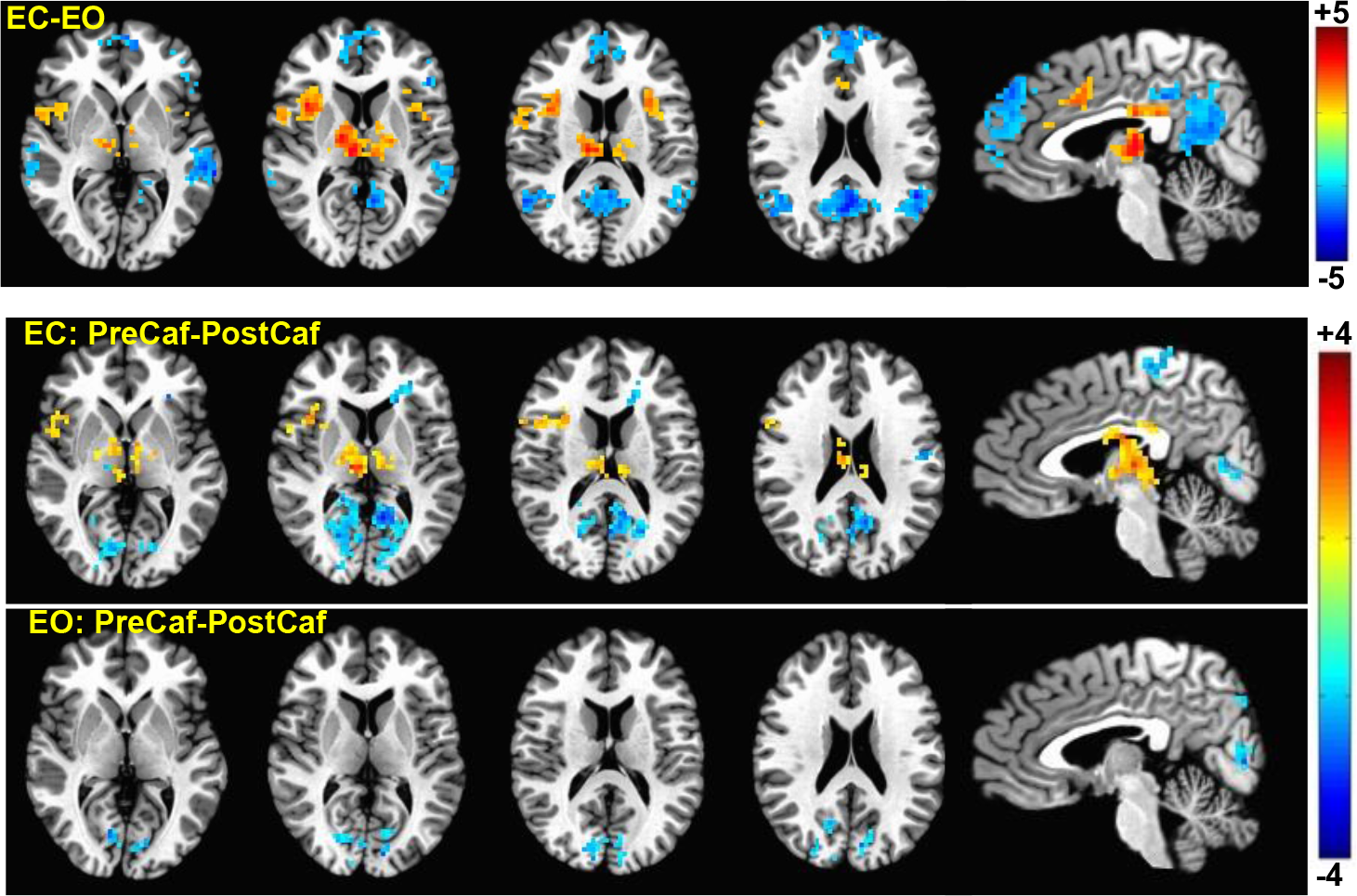
Group result z-scores (from 3ttestH—+) showing areas of significant difference between the vigilance correlation maps derived from different sessions. The top row shows the ECnonCaf minus EOnonCaf map (*p* < 0.05, corrected). The middle and the bottom rows display preCaf minus postCaf for eyes-closed and eyes-open conditions, respectively (*p* < 0.05 uncorrected).

### 5.2. Predictivity across conditions

Noting the differences in the vigilance correlation maps across different conditions, we examined the possibility of using the correlation map derived from one condition as a template for estimating the vigilance fluctuations occurring in another condition. The left panel of Figure 4 illustrates the predictivity results from 4 different scenarios:

1. ECnonCaf template predicting vigilance fluctuations in the ECnonCaf scans (EC → EC)
2. EOnonCaf template predicting vigilance fluctuations in the ECnonCaf scans (EO → EC)
3. EOnonCaf template predicting vigilance fluctuations in the EOnonCaf scans (EO → EO)
4. ECnonCaf template predicting vigilance fluctuations in the EOnonCaf scans (EC → EO)

**Figure 4:**
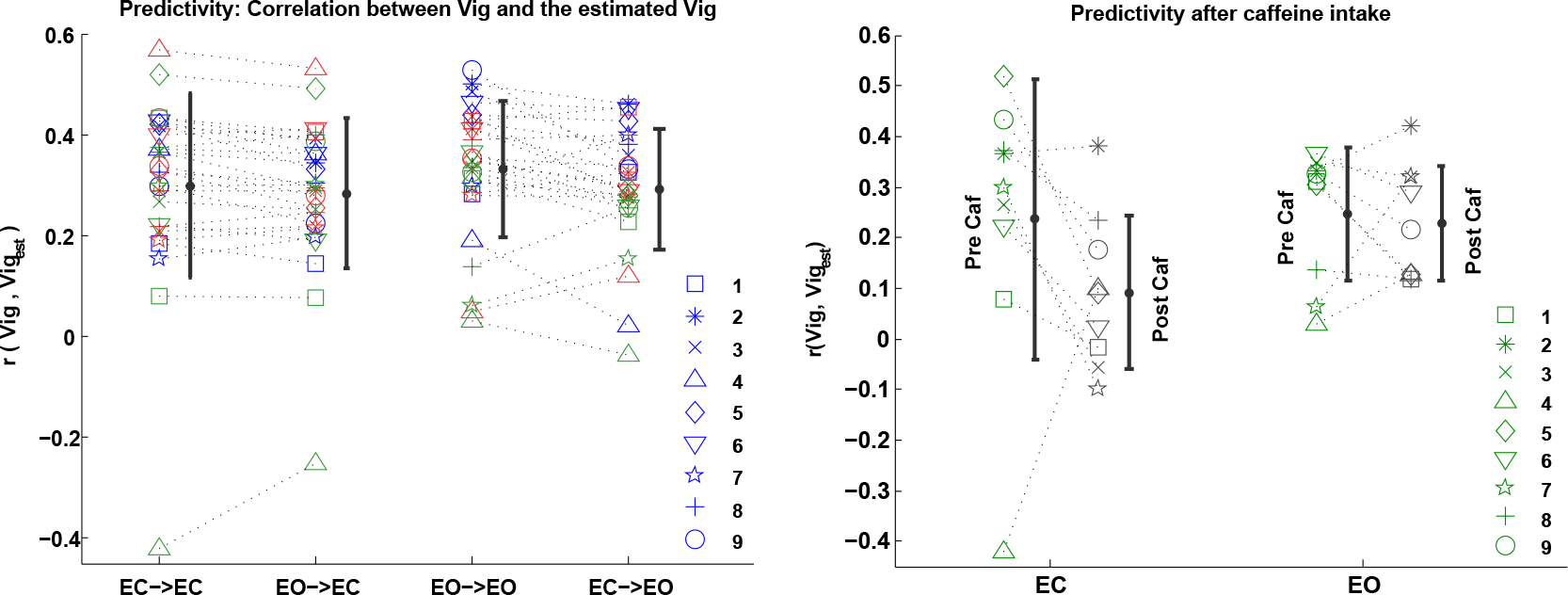
Predictivity: correlation between the EEG-based vigilance time series and the estimated vigilance using matching and non-matching templates across different conditions. Left panel: 1- “EC→EC”: ECnonCaf template predicting vigilance fluctuations of the ECnonCaf scans, 2- “EO→EC”: EOnonCaf template predicting vigilance fluctuations of the ECnonCaf scans, 3- “EO→EO”: EOnonCaf template predicting vigilance fluctuations of the EOnonCaf scans, and 4- “EC→EO”: ECnonCaf template predicting vigilance fluctuations of the EOnonCaf scans. Right panel: comparing the predictivity before and after caffeine. Each symbol represents a subject and different scanning sessions are shown with different colors.

The results from scenarios 1 and 3 are the same as in Figure 1 and are plotted again for comparison. Based on one-sample t-tests on the predictivity values when the templates were used across conditions, we found significant positive predictivity values in both scenarios 2 (EO→EC: *t*(26) = 9.2, *p* < 10^−6^, *d* = 1.7) and 4 (EC→EO: *t*(26) = 11.9, *p* < 10^−6^, *d* = 2.2).

Using paired t-tests we found that the EOnonCaf template can better predict the vigilance fluctuations of the EOnonCaf data as compared to the scenario where the ECnonCaf template was used on the same data (*t*(26) = 2.5, *p* = 0.01, *d* = 0.5). For the ECnonCaf data, no significant difference was found in the predictivity values when the ECnonCaf template was used as compared to when the EOnonCaf template was used (*t*(26) = 1.6, *p* = 0.1, effect size=0.3).

Predictivity for the caffeine scans was examined in both eyes-closed and eyes-open conditions using the average templates derived from the ECnonCaf and EOnonCaf scans, respectively. See the right panel of Figure 4. We found that in the EC scans, except for one outlier, the predictability was lower for the ECpostCaf scans as compared to ECpreCaf scans (*t*(8) = −1.3, *p* = 0.2; after outlier exclusion: *t*(7) = −3.6, *p* = 0.007, d=-1.3). In the EO scans, however, we did not find a significant difference between the EOpostCaf and EOpreCaf results (*t*(8) = −0.4, *p* = 0.6, *d* = −0.1). Using one-sample t-tests, we found that the overall predictivity after taking the caffeine reduced to a non-significant level in the ECpostCaf state (*t*(8) = 1.7, *p* = 0.1, *d* = 0.5), whereas for the EOpostCaf state it remained significant (*t*(8) = 5.9, *p* = 0.0003, *d* =1.9).

We also examined an additional scenario where the average templates derived from the post caffeine sessions were used to predict the vigilance fluctuations of the non-caffeine scans. We found a significant reduction in the predictivity values in the ECnonCaf state when the ECpostCaf template was used (*t*(26) = −2.3, *p* = 0.02, *d* = −0.4), whereas using the EOpostCaf template in the EOnonCaf condition did not have a significant effect on predictivity (*t*(26) = −1.1, *p* = 0.2, *d* = −0.2).

### 5.3. Predictivity and template similarity

For each scan, template similarity was defined as the spatial correlation between each individual template (*Temp_i_*) and the *Temp_LOO_* that was used for prediction. As shown in Figure 5, we found a significant relation between the predictivity and the template similarity values in both ECnonCaf (*r* = 0.93, *p* < 10^−6^) and EOnonCaf (*r* = 0.91, *p* < 10^−6^) conditions. In other words, the ability to predict vigilance fluctuations within an individual scan is strongly related to the spatial similarity between the individual template (i.e. correlation map between vigilance and BOLD for that scan) and the average LOO template that is used for the prediction process. An expression for the relation between the template similarities and the predictivity is presented in the appendix.

**Figure 5:**
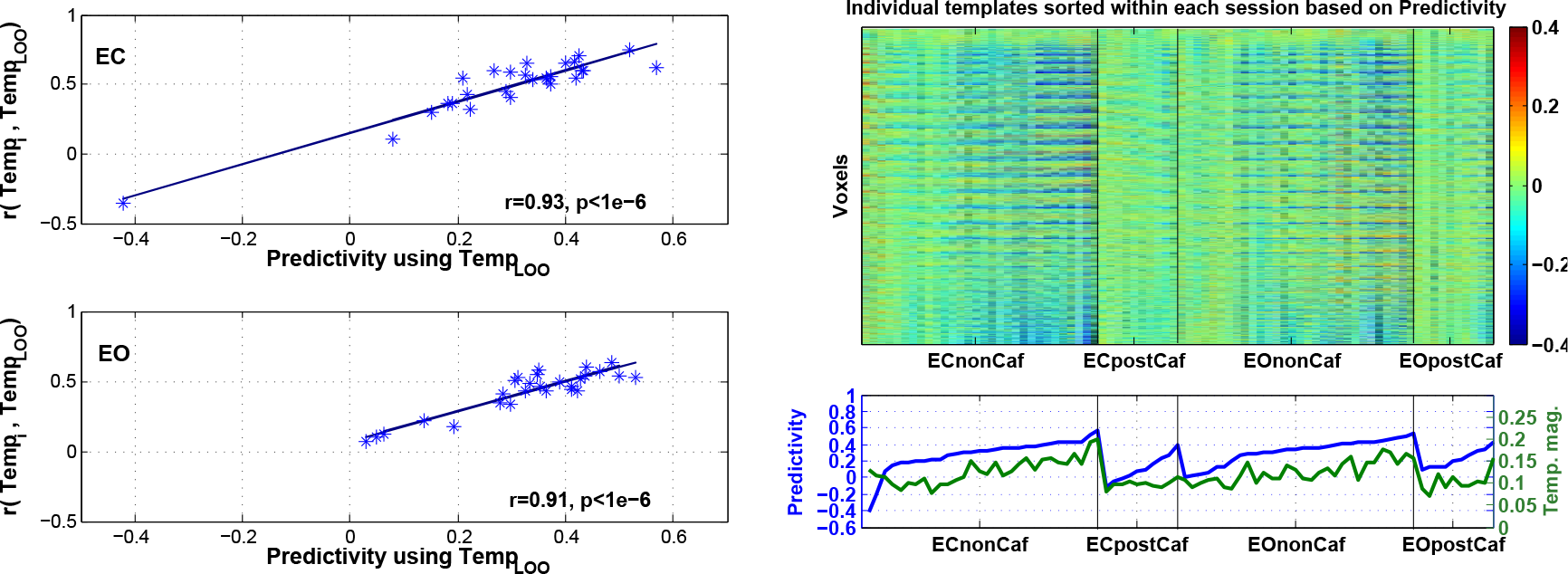
Left: Relation between predictivity and the similarity between *Temp_i_* and *Temp_LOO_* for ECnonCaf (top) and EOnonCaf (bottom) scans. Top right: Individual templates (*Temp_i_*) reshaped to 1-dimensional columns sorted based on the predictivity within each session, i.e. the subject on the left column in each session had the lowest predictivity and the subject on the rightmost column in each session had the highest predictivity. Bottom right: Predictivity values sorted within each session in blue and corresponding *Temp_i_* magnitude values across subjects in green.

For better visualization of the variability across the individual templates (*Temp_i_*), each template was reshaped to a 1-dimensional vector and then displayed as a column in the right panel of Figure 5. The templates (columns) were sorted within each session based on their predictivity values, i.e. the leftmost subject in each session has the lowest predictivity and the rightmost subject has the highest. The sorted predictivity values along with the template magnitude values (defined as the root mean square of each individual template) are shown in the second row. We found that there is a significant correlation between template magnitude and the predictivity values (*r* = 0.64, *p* < 10^−6^). In other words, the ability to predict the vigilance fluctuations within an individual scan is significantly related to the strength of the spatial map of correlations between the vigilance fluctuations and the BOLD time series. In the image of templates shown in the upper righthand portion of Figure 5, this is reflected as an increase in the visibility of distinct spatial patterns in the templates as predictivity increases (from left to right for each session). The template magnitude was also found to be related to global signal amplitude (*r* = 0.43, *p* = 4 × 10^−5^), with larger template magnitudes corresponding to larger global signal amplitudes. Template magnitudes were reduced after caffeine in both eyes-closed (postCaf-preCaf: *t*(8) = –2.6, *p* = 0.03, *d* = −0.86) and eyes-open (postCaf-preCaf: *t*(8) = –2.4, *p* = 0.04, *d* = –0.8), reflecting a decrease in the strength of the spatial patterns in the individual templates.

### 5.4. Vigilance amplitude

As a measure of the variations in vigilance levels within a scan, we defined vigilance amplitude (*aVig*) as the standard deviation of the vigilance time series. The first row of Figure 6 illustrates the relation between the amplitude of the estimated vigilance (*aVig_est_*) and the amplitude of the EEG-based vigilance time series (*aVig*). The amplitude of the estimated vigilance was found to be significantly correlated with the amplitude of the EEG-based vigilance in both ECnonCaf (*r* = 0.63, *p* = 0.0005) and EOnonCaf (*r* = 0.54, *p* = 0.003) conditions. We also found that the predictivity is significantly related to the amplitude of the estimated vigilance in both ECnonCaf (*r* = 0.63, *p* = 0.0004) and EOnonCaf (*r* = 0.47, *p* = 0.01), with higher values of vigilance amplitude associated with higher predictivity.

**Figure 6:**
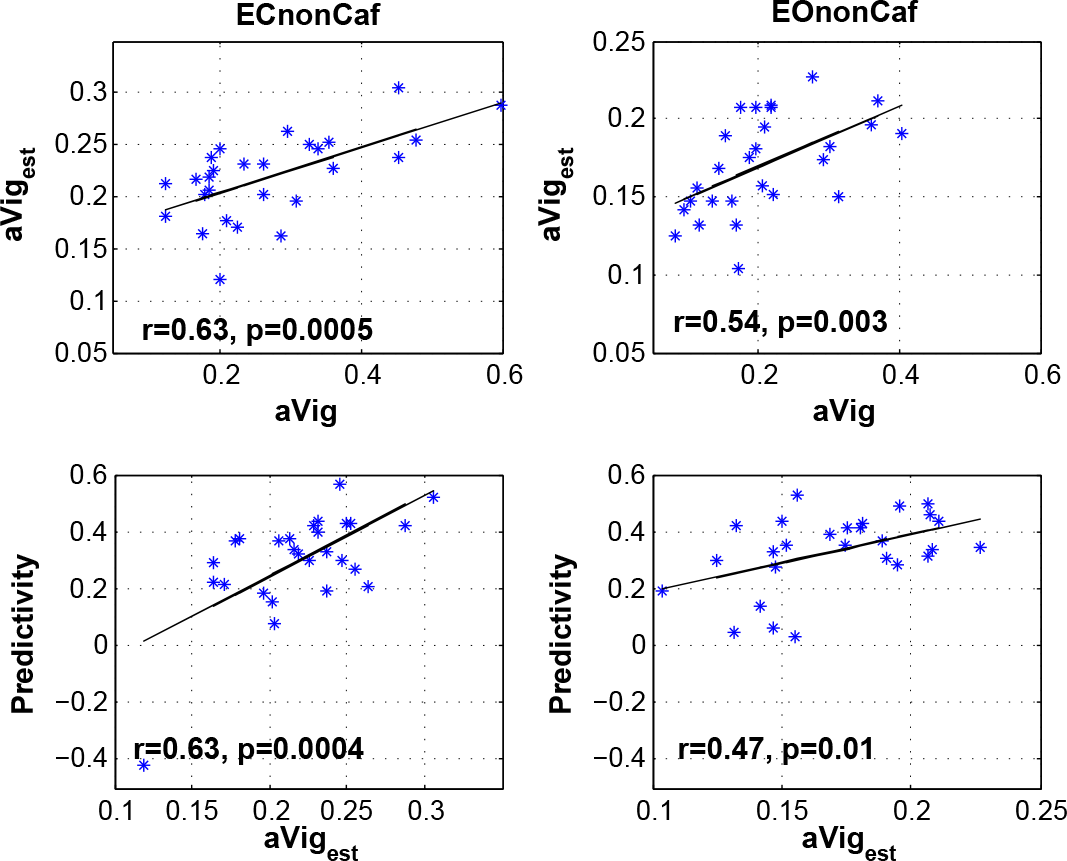
Top row: Relation between the amplitude of the estimated vigilance time series (*aVig_est_*) and the amplitude of the EEG-based vigilance time series (*aVig*) with the linear fit shown by the black lines. Bottom row: Relation between predictivity and the amplitude of the estimated vigilance time series (*aVig_est_*) with the linear fit shown by the black lines.

As shown in Figure S1, we found that the amplitude of vigilance is correlated with the template similarity measures (ECnonCaf: *r* = 0.4, *p* = 0.03, EOnonCaf: *r* = 0.66, *p* = 0.0002), such that a higher vigilance amplitude corresponds to a greater level of similarity between a subject’s individual template and the template used for predctivity, i.e. *Temp_LOO_*. In addition, changes (postCaf - preCaf) in vigilance amplitude after caffeine intake were significantly related to changes in the similarity between the templates (EC: *r* = 0.86, *p* = 0.002, EO: *r* = 0.6, *p* = 0.07).

### 5.5. Relation between vigilance, GS and predictivity

A negative correlation between the vigilance time series and fMRI global signal was reported in our prior work (Falahpour et al., 2016) and is plotted in Figure S2 for the reader’s convenience. Predictivity using the template approach is compared against the correlation between global signal and vigilance in Figure 7. Using a paired t-test we found that using template for vigilance estimation is significantly better than using global signal as an estimator in both the ECnonCaf (*t*(26) = 3.4, *p* = 0.002, *d* = 0.6) and EOnonCaf (*t*(26) = 5.5, *p* = 9 × 10^−6^, *d* = 1.06) conditions.

**Figure 7:**
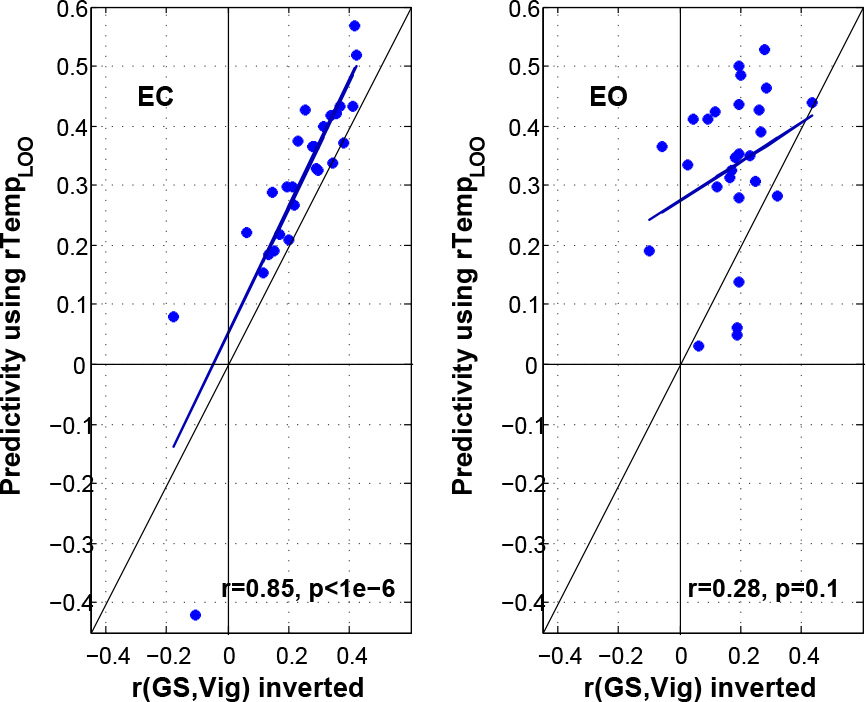
Predictivity using template versus the correlation between vigilance and global signal (inverted) for ECnonCaf and EOnonCaf conditions. The linear fits are shown by dark blue lines. The vertical, horizontal, and slanted black lines represent indicate the x-axis, y-axis, and line of unity, respectively.

## 6. Discussion

In this paper, we have shown that vigilance fluctuations in humans during a typical resting state fMRI scan, with either eyes-open or eyes-closed, can be estimated from the fMRI data alone using the template-based approach. Fluctuations in the power of different EEG frequency bands are known to be related to vigilance or alertness as decreases in vigilance are characterized by increases in delta and theta band power and decreases in alpha band power (Klimesch, 1999). Prior studies have shown that EEG alpha power is negatively correlated with fMRI signals in widespread regions of the brain, including the visual and fronto-parietal cortices, and is positively correlated in the thalamus (Goldman et al., 2002; Laufs et al., 2003; Sadaghiani et al., 2010). Here we used the ratio of the EEG power in the alpha band to the power in the theta and delta bands in order to derive fMRI template maps that can be used for estimating vigilance fluctuations in the absence of EEG. Areas in our template showing strong correlations with this EEG vigilance measure are in agreement with those identified in prior studies (Goldman et al., 2002; Laufs et al., 2003).

The approach proposed in (Chang et al., 2016) was applied here, but there were several differences between that study and the current work that are worth discussing: 1) the original work evaluated the performance of the proposed method using data acquired from 4 monkeys sitting in near complete darkness, while our work provides the first application of this approach to human data. We found that the predictivity values, defined as the correlation between the estimated vigilance (or arousal) and the EEG-based vigilance time series, are comparable between the two studies (i.e. with mean=0.4, median=0.39 and interquartile range (IQR)=0.15 for 14 runs from 2 monkeys with LFP recordings, and mean=0.56, median=0.56 and IQR=0.19 for 33 runs from 4 monkeys with eye signal in the original work; and with mean=0.31, median=0.34 and IQR=0.15 for 54 runs in this study). 2) In the prior work, the monkeys’ eye behavior was unconstrained and relatively long scan durations were used (each scan was about 30 minutes), such that the monkeys were more likely to experience large fluctuations in vigilance and wakefulness over the course of the scan. In contrast, here we evaluated the performance of the approach using a shorter scan duration (5 minutes) and conditions (e.g. eyes open fixation or eyes closed) that are more typical of human resting-state fMRI protocols. 3) Our results indicated that there is a wide range of variations in the predictivity values across subjects and conditions, a feature that was not examined in the original study.

Variability in the performance of the vigilance estimator appeared to be related to the amplitude of the vigilance fluctuations experienced by the subject within a given scan. In addition, we found that the amplitude of vigilance fluctuations seems to play a role in forming the structure of an individual’s template, where higher amplitudes correspond to more pronounced structures in the spatial maps and greater similarity with the average template. These observations indicate that a greater variation in vigilance levels across the scan is accompanied by stronger modulation of vigilance-related brain activity and, in turn, a stronger match between the resulting fMRI data and a general template of vigilance-related fMRI activity. Conversely, when the variability in the vigilance levels is small, there is a reduced modulation of brain activity, making it difficult to estimate the vigilance fluctuations. As an example, we found that the predictivity decreases significantly after caffeine consumption in the EC condition, with a corresponding reduction in vigilance amplitude. This might be a limiting factor in applying the method on subjects with a relatively constant level of alertness (e.g. being very vigilant or caffeinated). However, if the vigilance state is indeed relatively constant across a scan, there is also less of a need to estimate vigilance fluctuations in the first place, as the impact of vigilance on that particular dataset would be minimal.

In general, it is likely that most resting-state studies will find that there is broad range in the strength of vigilance fluctuations across different scans (e.g. between subjects and conditions), such that the ability to predict the vigilance fluctuations will also vary across scans in a manner similar to what we found in this study. In the analysis of these studies, it may turn out that there is not a great need to actually estimate the time course of vigilance fluctuations. Instead, it may be sufficient to estimate the amplitude of the vigilance fluctuations, as such a measure could be used as a covariate in the analysis process. Our results indicate that the template-based approach can offer reasonable estimates of vigilance amplitude.

Prior studies have shown that functional brain connectivity changes dynamically (Hutchison et al., 2013) and that one influential factor is the vigilance state (Haimovici et al., 2017; Chang et al., 2013). Given that fluctuations in a subject’s wakefulness state are likely to occur during a typical resting state fMRI experiment (Tagliazucchi and Laufs, 2014), an estimate of vigilance can be helpful in the interpretation of experimental findings. In particular, when analyzing the data from different cohorts of subjects, one major assumption is that the subjects are in similar states of wakefulness, both within and between groups. When this assumption is violated, inter-subject and inter-group differences in vigilance can significantly alter estimates of functional connectivity differences. Our work suggests that template-based vigilance estimates can be used to account for inter-subject and inter-group vigilance differences in the analysis and interpretation of the data.

Here, vigilance estimation was also examined in different scenarios: 1) using the ECnonCaf template on EOnonCaf data, and vice versa; 2) using the ECnonCaf and EOnonCaf templates on ECpostCaf and EOpostCaf data, respectively; and 3) using the caffeine templates, i.e. ECpostCaf and EOpostCaf on non-caffeine data, i.e. ECnonCaf and EOnonCaf, respectively. Our results indicated that even though there are significant differences between the ECnonCaf and EOnonCaf templates, they can still be used interchangeably. In terms of the performance, however, the EOnonCaf template seems to outperform the ECnonCaf template.

Interestingly, the differences between the ECnonCaf and EOnonCaf templates were localized to regions forming the default mode network (DMN), along with the insula (which is part of the salience network; (Seeley et al., 2007)) and thalamus. A positive correlation between DMN BOLD signals and visual alpha power was reported in a prior study (Mo et al., 2013) when comparing EC to EO (EO minus EC). While their EO-EC contrast maps are similar to our results (considering the difference in polarity as we show EC minus EO), their correlation maps obtained in each condition differ from our findings as they found positive correlations between DMN BOLD and visual alpha only in the EO condition, and no significant correlations in EC. In contrast, we found that there are negative correlations between the BOLD signal from DMN regions and vigilance fluctuations in the EC state, and non significant correlations in the EO condition. This could be due to the differences in the fMRI preprocessing methodologies as they removed global signal components from their data. In addition, their use of visual alpha power differs from our use of an EEG vigilance measure, which considers the ratio between the power in the alpha band and the power in the combined delta and theta bands.

The DMN is known to be active during mind wandering and day dreaming, and to deactivate during many external goal-oriented tasks (Buckner et al., 2008). Additionally, Braboszcz and Delorme (2011) found that, during mind wandering in the EC condition, the EEG activity in low frequency bands (delta and theta) increased whereas the activity in the high frequencies (alpha and beta) decreased. This implies that during lower vigilance states subjects experience more mind wandering and day dreaming with a corresponding increase in DMN activity. In agreement with this prior work, our findings of a negative correlation in EC are consistent with a picture in which higher vigilance states may be associated with higher levels of external orientation (and lower DMN activity), whereas lower vigilance states may be associated with higher levels of self-referential activity (and higher DMN activity). This relation is found in the EC condition where the amplitude of vigilance fluctuations is also higher (EC-EO: *t*(26) = 3.9, *p* = 0.0004, *d* = 0.76), likely because it is a condition more conducive to periods of drowsiness.

After taking caffeine, the predictivity became insignificant only in the EC condition. This is expected if one compares the ECpostCaf templates with the other templates derived from non caffeine sessions, both at the individual level displayed on the right panel of Figure 5 and at the group level shown in Figure 2. The correlation pattern in the brain that is visible in the other templates is greatly diminished in the ECpostCaf condition and consequently the predictivity values are significantly reduced when the ECnonCaf template is used to estimate the vigilance fluctuations of the post caffeine session. We quantified the reduction in visibility by showing that the template magnitudes were significantly reduced after the caffeine intake. In addition, the templates magnitudes were found to be significantly correlated with both the predictivity and the amplitude of the global signal. Our findings are roughly consistent with prior studies showing that caffeine reduces the standard deviation of the additive global signal, with a stronger reduction in the EC condition (Wong et al., 2012, 2013). Given the correlation between the global signal and vigilance fluctuations reported both here and in our preliminary work (Falahpour et al., 2016), caffeine-induced reductions in both global signal and vigilance fluctuations (observed in 6 out of 9 subjects) are to be expected. It is likely that caffeine induces a more constant vigilance state, and therefore the reduction in predictivty reflects two related effects: (1) vigilance fluctuations are decreased and hence (2) fMRI signals related to vigilance modulation are also decreased. Both of these factors can reduce the efficacy of the template-based approach.

Although the interpretation and the origins of the global signal are not well understood, there is evidence that the global signal contains contributions from neural sources (Wong et al., 2013; Liu et al., 2015; Schölvinck et al., 2010). For example, our earlier work demonstrated that there is a significant correlation between the global signal and vigilance fluctuations (Falahpour et al., 2016). Therefore, it is possible that the global signal may serve as an estimator of a subject’s vigilance variation during rest. Here, we examined this possibility directly and demonstrated that using the joint spatial information provided in a template pattern can improve the estimation of vigilance beyond that obtained with the global signal, consistent with the findings of (Chang et al., 2016). In particular, the improvement gained by using the template was higher in the EO condition as compared to the EC condition; in the EC condition, the improvement was significant but the effect size was not large. This might be explained by the fact that in the EC condition, which is more conducive to drowsiness, the global signal may be more dominated by large-scale neural activity related to vigilance changes as compared to other neural and non-neural fluctuations (e.g., head motion and physiological noise). Therefore, the global signal might provide a reasonable approximation of vigilance during the EC condition when the subject are likely to be more drowsy, but one must be cautious given the fact that global signal reflects neural activity of interest from distributed brain networks.

In conclusion, our findings suggest the feasibility of using a template-based approach to estimate vigilance fluctuations directly from resting-state fMRI in human subjects. Such an fMRI data-driven estimate may help to increase the sensitivity of resting-state fMRI data to neural effects of interest, particularly when external vigilance monitoring (such as with EEG) is not feasible. Our results also indicate that there might be a dynamic relationship between DMN activation and vigilance state, but further work is required to investigate this observation in more detail.

## 7. Acknowledgements

This work was supported in part by NIH grant R21MH112155.

## Appendix A. Predictivity and template similarity

Here we show the relation between predictivity and the spatial similarity between an individual’s template and the LOO template used for estimation. To provide insight, we first consider a simplified estimator based on the inner products of the BOLD data with a LOO template. We then consider normalization of the simplified estimator, which yields the correlation coefficient based estimate that is used in the paper.

### Appendix A.1. Using the inner product

Let *V_i_* and *X_i_* denote the EEG-based vigilance time series and BOLD fMRI data from subject i. We can write them as:

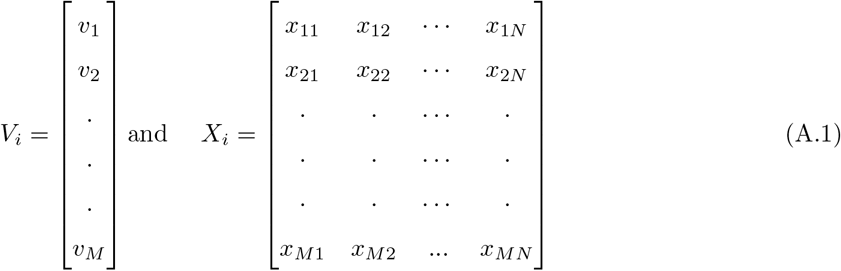

where *M* and *N* are the number of time points and the number of voxels in the brain, respectively. For each subject, we can define the template (*T_i_*) as the inner product between vigilance and fMRI data:

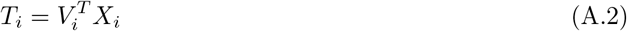

To estimate the vigilance time series for the *j*th subject, we first form an LOO template as follows:

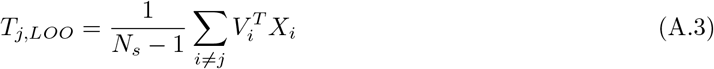

where *N_s_* is the number of subjects. Then for each subject, the vigilance time series may be estimated by computing the inner products between the LOO template and BOLD data:

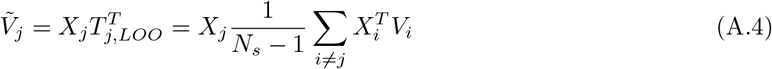

As a measure of similarity, we can use the inner product between the estimated vigilance 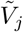 and the EEG-based vigilance *V_j_*, which is given by:

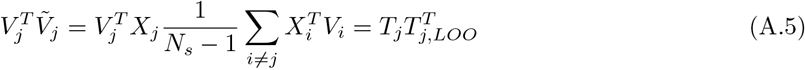

Therefore, if the inner product is used, the similarity between the estimated vigilance and the EEG-based vigilance for each scan is equal to the similarity between the scan’s template and the LOO template 490 used for estimation.

### Appendix A.2. Using the correlation coefficient

We define *σ_X_i__* as the 1 × *N* row vector consisting of the standard deviations of the voxel time series in *X_i_* and *s_X_i__* as the 1 × *M* row vector consisting of the spatial standard deviations of the images in *X_i_* at each time point, such that

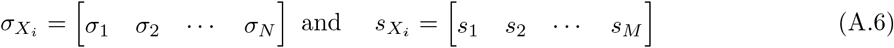

Without loss of generality we assume that the mean of all vectors are removed prior to calculating the correlations. We define the template (*T_i_*) as the vector of correlation coefficients between the vigilance time series and fMRI data, such that

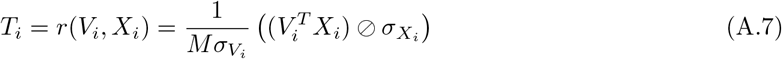

where ∅ denotes element-wise division. Similar to the previous section, we form the LOO template and estimate the vigilance as follows:

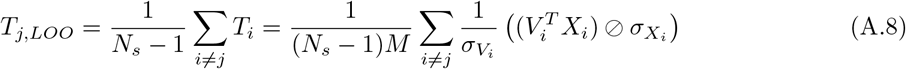

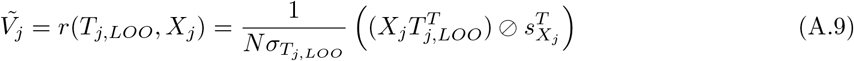

where *σ_T_j, LOO__* is the standard deviation of the *j^th^* LOO template. The correlation coefficient between the estimated vigilance and the EEG-based vigilance time series can be written as:

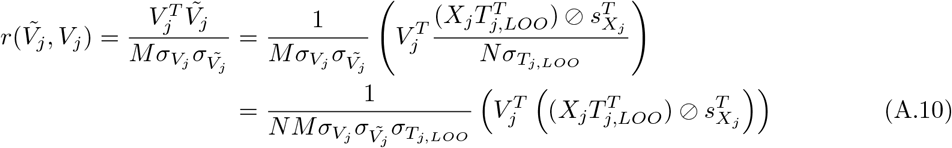

where *σ_V_j__* and 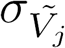 are the standard deviations of the vigilance and estimated vigilance time series, respectively.

The similarity between each subject’s individual template and the average LOO template can be written as:

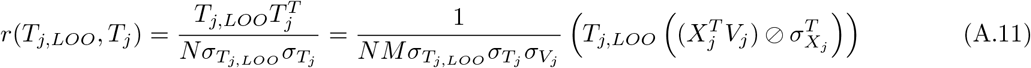

where *σ_T_j__* is the standard deviation of the individual template.

The relation between the two above equations can be simplified to:

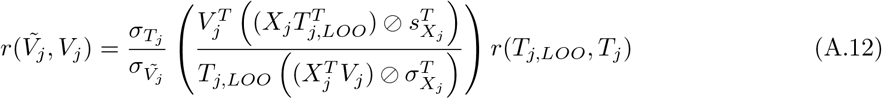

Thus, for each scan, the correlation coefficient between the estimated vigilance and the EEG-based vigilance is equal to a scaled version of the correlation coefficient between the individual’s template and the LOO template used for estimation. The scaling factor is equal to 1 if all the standard deviations terms are equal to 1. In practice the scaling factors are distributed around 1 as shown by the distribution of points around the line of unity in Figure S3 with a mean slope of 0.82.

## Supplementary figures

**Figure S1:**
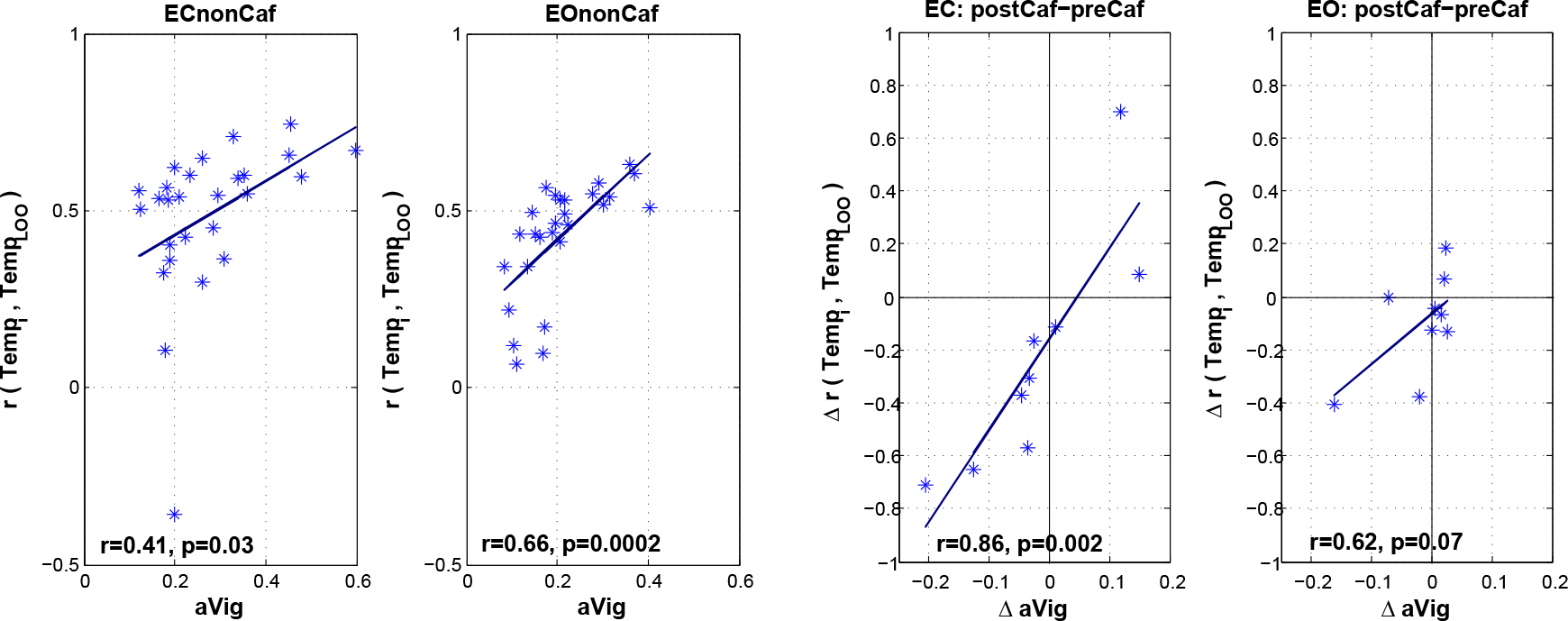
Left: Similarity between the individual templates (*Temp_i_*) and *Temp_LOO_* in non caffeine sessions versus the amplitude of vigilance time series (*aVig*). Right: Changes (post-caffeine minus pre-caffeine) in the correlation between *Temp_i_* and *Temp_LOO_* derived from non caffeine sessions versus the changes in the vigilance amplitude after taking caffeine. The linear fits are shown by dark blue lines. The vertical and horizontal black lines show zeros for x and y axes.

**Figure S2:**
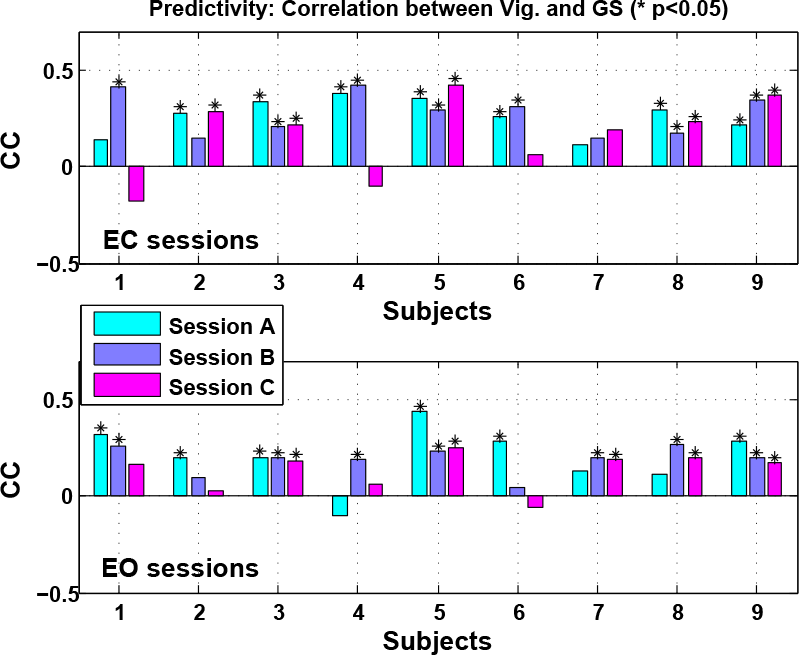
Correlation between the EEG-based vigilance time series and global signal (inverted) for ECnonCaf in the top row and EOnonCaf in the bottom row

**Figure S3:**
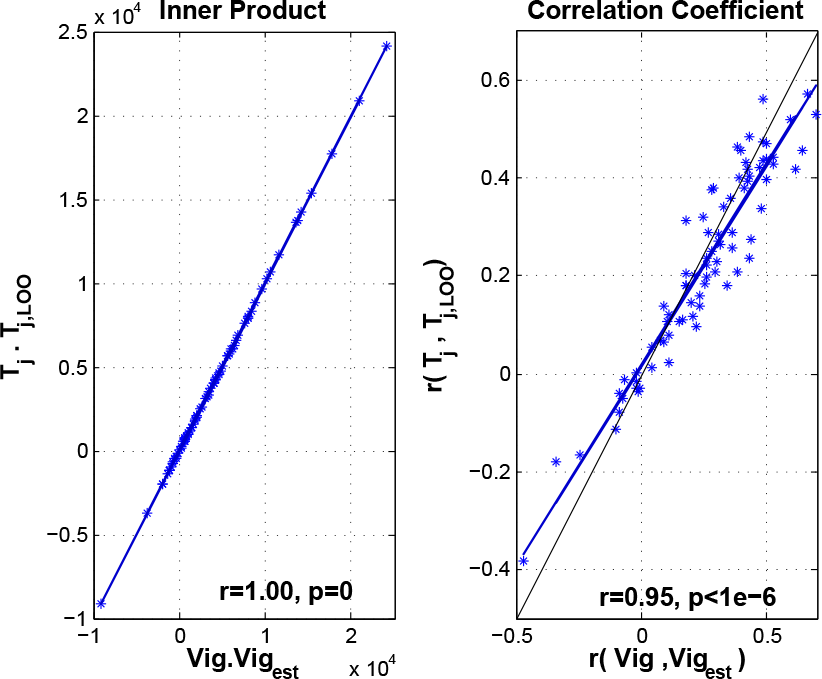
Left: Inner product of the individual template (*Temp_i_*) and *Temp_LOO_* versus the inner product between the EEG-based vigilance time series and the estimated vigilance. Right: Correlations between *Temp_i_* and *Temp_LOO_* vs the correlation between the EEG-based vigilance time series and the estimated vigilance. The blue and black lines show the linear fits and the unity lines, respectively. The slopes for the linear fits are 1 and 0.82 for the inner product on the left and the correlation on the right panel, respectively.

